# Catalytic and Structural Insights into Neil3-Dependent Unhooking of Endogenous Abasic DNA Crosslink

**DOI:** 10.64898/2026.01.19.700286

**Authors:** Andrea Huskova, Barbora Landova, Viktoria Benova, Martin Klima, Kamil Hercik, Evzen Boura, Jan Silhan

## Abstract

Abasic (Ap) sites arise frequently in genomic DNA and can form interstrand crosslinks (Ap-ICLs) that block DNA replication and threaten genome stability. The DNA glycosylase NEIL3 is required for replication-coupled repair of Ap-ICLs, yet its catalytic mechanism has remained unclear, as biochemical studies report lyase-dependent strand cleavage whereas cellular systems indicate incision-free unhooking.

Here, we show that the catalytic outcome of NEIL3 is determined by the N-terminal processing of its NEI domain. Using biochemically and structurally defined NEI variants, we demonstrate that a native-like processed form (V2M), in which valine 2 is replaced by an initiating methionine, efficiently unhooks Ap-ICLs by releasing the crosslinked strand without generating toxic DNA strand breaks, and without β- or δ-elimination. In contrast, an unprocessed form (M1) exhibits elevated Ap-lyase activity and generates strand breaks. Time-resolved Schiff-base trapping in the presence of a reducing agent reveals distinct high-molecular-weight intermediates during Ap-ICL unhooking. A crystal structure of NEIL3 bound to native-like substrate in form of a single-stranded DNA identifies features underlying its preference for fork-like substrates.

Together, these findings reconcile previously conflicting models of NEIL3 function and define a mechanistic framework for replication-coupled repair of endogenous crosslinks, Ap-ICL, preserving fork integrity.

## Introduction

DNA is intrinsically unstable under physiological conditions. Spontaneous hydrolytic reactions lead to frequent base loss, generating abasic (Ap) sites at rates estimated to reach thousands of events per cell per day (1–3). These lesions are central intermediates of base excision repair (BER) but are themselves chemically reactive and prone to further transformation if not promptly resolved (4–6). Accumulation or misprocessing of Ap sites can therefore contribute to genome instability and cell death (4).

One chemically distinct fate of Ap sites is the formation of interstrand DNA crosslinks (ICLs). The ring-opened aldehyde form of an abasic sugar can react with nucleophilic sites on the opposing strand, most prominently adenine, resulting in covalent interstrand linkages (7, 8). These Ap-ICLs form under physiological conditions, are chemically stable within duplex DNA, and are unavoidable endogenous lesions (9–11). Because they physically tether the two DNA strands, Ap-ICLs constitute absolute blocks to DNA replication and transcription and must be actively removed to preserve cell viability (4, 12, 13).

ICLs are among the most cytotoxic DNA lesions, and their repair has been studied extensively in the context of chemically induced crosslinks, such as those generated by platinum compounds or nitrogen mustards (4, 12, 14). In eukaryotic cells, repair of these lesions is tightly coupled to DNA replication and orchestrated by the Fanconi anaemia (FA) pathway, which coordinates nucleolytic incisions, translesion synthesis, and homologous recombination (15, 16). Although effective, this process involves the formation of DNA double-strand breaks (DSBs) and is therefore intrinsically mutagenic. The existence of endogenous Ap-ICLs raises the question of whether cells employ specialised repair mechanisms that minimise secondary DNA damage when dealing with these chemically simpler, yet frequent, lesions.

Seminal work has identified the DNA glycosylase NEIL3 (Nei-like glycosylase 3) as a key factor required for replication-coupled repair of Ap-ICL(**17**) (17). Using *Xenopus* egg extracts, Semlow et al. demonstrated that NEIL3 promotes unhooking of Ap-ICLs at stalled replication forks without generating detectable DSBs (17). In this pathway, NEIL3-dependent repair allows replication to resume without activation of the FA machinery, whereas loss of NEIL3 function reroutes Ap-ICLs to canonical FA-dependent processing (18). Recruitment of NEIL3 to stalled forks is regulated by TRAIP-mediated ubiquitylation of the CMG helicase, positioning NEIL3 at the leading edge of replication stress (19).

NEIL3 belongs to the Fpg/Nei family of DNA glycosylases, which typically function as bifunctional enzymes coupling base excision to β- and δ-elimination reactions that cleave the DNA backbone at abasic sites (20, 21). Structural and mechanistic studies of bacterial Fpg/Nei homologues have established a conserved catalytic mechanism initiated by an N-terminal proline, leading to strand incision and formation of blocked termini (22–25). In contrast to these enzymes, NEIL3 exhibits a strong preference for single-stranded and forked DNA substrates and displays limited activity on fully duplex DNA (10, 26–29). Structural analysis of the mouse NEIL3 NEI domain revealed an architecture compatible with ssDNA engagement, consistent with its specialised function at replication forks (26).

Despite this apparent specialisation, purified NEIL3 has been reported to possess Ap lyase activity *in vitro*, generating strand breaks at abasic sites and crosslinks (10, 27, 30). This observation stands in apparent contradiction to cellular and extract-based studies indicating that NEIL3 resolves Ap-ICLs without inducing DNA strand breaks (17, 19). Reconciling these conflicting outcomes has remained a central unresolved question in understanding NEIL3-dependent repair.

One potentially overlooked factor is the catalytic state of the NEIL3 N-terminus. In contrast to canonical Fpg/Nei enzymes, NEIL3 lacks a conserved N-terminal proline and instead initiates with a valine residue whose removal depends on N-terminal methionine excision during protein expression (30, 31). In *Escherichia coli*, the efficiency of methionine removal is governed by the identity of the penultimate amino acid, leading to heterogeneous populations of processed and unprocessed proteins (32–34). For NEIL3, incomplete N-terminal processing has been reported during recombinant expression and purification, raising the possibility that distinct catalytic behaviours observed *in vitro* may reflect differences in N-terminal identity rather than intrinsic mechanistic plasticity (31). Previous studies have proposed that NEIL3 activity may be modulated by its auxiliary domains or by autoinhibitory interactions involving zinc finger motifs (35, 36). However, the contribution of N-terminal processing of the NEI domain itself to catalytic outcome has not been systematically addressed. Given the central role of the N-terminus in Fpg/Nei-mediated chemistry, even subtle changes in N-terminal identity could profoundly affect the balance between glycosylase-only activity and strand-cleaving lyase reactions.

Here, we investigate the molecular basis of Ap-ICL unhooking by NEIL3 using biochemically defined variants of the NEI domain that differ solely in their N-terminal processing state. To isolate the contribution of N-terminal identity, we compared an unprocessed methionine-initiated form (M1) with a native-like processed variant (V2M), which lacks Met1 and serves as a substitute for the valine-initiated enzyme because catalysis depends on the N-terminal amine. This strategy eliminates ambiguity arising from incomplete post-translational processing and enables direct attribution of catalytic outcome to N-terminal identity. Using defined forked DNA substrates containing site-specific abasic lesions, we show that the processed variant efficiently unhooks Ap-ICLs without inducing β- or δ-elimination, whereas the unprocessed form preferentially promotes strand incision via lyase activity. These findings reconcile previously conflicting models of NEIL3 function and define a mechanistic framework for replication-coupled repair of endogenous DNA crosslinks that preserves replication fork integrity.

## Materials and Methods

### Mutagenesis, expression and purification of mNeil3 (M1 and V2M forms)

The NEI domain of mouse Neil3 (mNeil3), with valine (V2M) as the first amino acid, was generated by site-directed mutagenesis using PCR, in which the initial methionine codon (M1) was deleted, and the first codon for valine serves as alternative initiation codon for methionine in this position (32–34). This modification was introduced into a previously published construct encoding the mNEI domain with a C-terminal His-tag, cloned in the pET-28 vector (35). The primers used for this mutagenesis were: PrimerF 5′-GTG GAA GGG CCA GGG TGT ACA C-3′ and PrimerR: 5′-GTA TTT ATC TCC TTC TTA AAG TTA AAC AAA ATT ATT T-3′.

### Expression and purification of His-tagged mNeil3 (M1 and V2M forms)

His-tagged mNeil3 proteins (M1, V2M and K82A variants) were expressed in *E. coli* NiCo21 (DE3) cells. Protein expression was carried out as previously described using standard laboratory protocols (35, 37, 38). Briefly, bacterial pellets were resuspended in lysis buffer containing 20 mM Tris pH 8.0, 300 mM NaCl, 20 mM imidazole pH 8.0, 2 mM β-mercaptoethanol (β-ME), and 10% glycerol, and lysed by sonication. Cell debris was removed by centrifugation, and the resulting supernatant was subjected to immobilised metal affinity chromatography on an Ni-NTA resin by 30-minute incubation at 4 °C on a rotary tube roller. After incubation, the resin was extensively washed with lysis buffer, and bound proteins were eluted using elution buffer containing 20 mM Tris pH 8.0, 300 mM NaCl, 300 mM imidazole pH 8.0, 2 mM β-ME, and 10% glycerol.

The eluate was desalted on HiPrep 26/10 Desalting column (Cytiva) and loaded onto a cation exchange HiTrap SpSepharose (Cytiva) equilibrated with buffer A (70 mM Tris pH 8.0, 125 mM NaCl, 10% glycerol, 2 mM β-mercaptoethanol (β-ME)). Bound proteins were further purified using a salt gradient on an ÄKTA purifier system (Cytiva). To remove the His-tag, the eluted protein was digested with 3C protease (prepared in-house) during overnight dialysis against gel filtration buffer (20 mM Tris pH 8.0, 200 mM NaCl, 2 mM β-ME, 10% glycerol) followed by gel filtration on chromatography Superdex 75 increase 10/300GL (Cytiva) in the same buffer. The purified protein was concentrated to 10 mg/mL and stored frozen for further use.

### Crystallisation of M1 NEI and V2M NEI Domains of Mouse Neil3

Trapped protein–DNA complexes were prepared as described previously (35). In brief, M1 and V2M NEI domains of mouse Neil3 were each incubated with DNA substrate containing the lesion TCCA[U]GTCT (Eurofins), in a 1:1.25 molar ratio (200 μM protein: 250 μM DNA) in a buffer containing 20 mM Tris pH 8.0, 125 mM NaCl, and 0.5 mM TCEP. The reaction was initiated by the addition of 50 mM sodium cyanoborohydride (Na[BH₃(CN)]) and monitored by SDS–PAGE. After completion, excess Na[BH₃(CN)] was removed via a 5 ml desalting column (Cytiva), and the trapped complex was purified using a Q HP anion exchange column (Cytiva).

Crystallisation trials were carried out using the sitting-drop vapour diffusion method at 18 °C, with drops set up on a Mosquito crystallisation robot (TTP Labtech, Melbourn, UK). Commercial screens, including ProPlex, JCSG Core I & II, Helix, and Morpheus (Molecular Dimensions, Sheffield, UK), were used, with drop ratios of 200 nl protein to 200 nl reservoir solution. Crystals typically appeared within 48 hours. Suitable crystals for diffraction were obtained in a condition containing 50 mM HEPES pH 7.0, 5 mM MgCl₂, and 25% PEG MME 550. Crystals were cryoprotected by soaking in mother liquor supplemented with 20% glycerol and flash-frozen in liquid nitrogen. Initial diffraction testing was performed on a home-source X-ray generator (Cu-Kα). The final dataset, resulting in structure determination (PDB ID: **8B9N**), was collected at BESSY II synchrotron (Helmholtz-Zentrum Berlin, Germany).

### Ap-ICL crosslink formation and purification

The reaction buffer was optimised to its final composition of 20 mM Tris pH 7.4, 140 mM NaCl, 0.5 mM TCEP, and 5% glycerol. Experiments were conducted under physiological conditions. Labelled DNA oligonucleotides (Supplementary Figure 2) were mixed with unlabelled complementary oligonucleotides in a ratio of 1:1 and annealed in a Biometra T professional Thermocycler by heating at 95°C for 5 minutes and left to cool to room temperature. The final concentration in the reaction buffer of dsDNA was 40 μM. Reactions were carried out in a 50 μl volume. The Ap site was generated by adding 1 ul Uracil-DNA-glycosylase (UDG) (New England Biolabs) per reaction and kept for 5 min at room temperature. Subsequently, UDG was deactivated by heating the reaction in another round of annealing in the thermocycler. Reactions were kept at 37°C. Crosslinked DNA (Ap-ICL) was isolated from polyacrylamide urea gel (TBE, 15% acrylamide/bisacrylamide (19:1) and 8M urea). Separated bands were visualised using HeroLabUVT-20 S/M/L, and the crosslinked band was cut out and transferred to a tube containing 200 μl of elution buffer composed of 20 mM Tris pH 7.4, 140 mM NaCl, 0.5 mM TCEP, 5% glycerol. The tube was left in a rotating mixer at 8°C overnight. The sample was centrifuged at 800 rpm for 1 min, and the supernatant was transferred into desalting columns (Cytiva MicroSpin Columns G-25) equilibrated with an elution buffer to remove excess urea.

### Ap and Ap-ICL Enzymatic Assays and Characterisation of 3′-product

Enzymatic assays to determine cleavage products from the Ap lyase activity of mouse Neil3 were performed in 20 µl reaction buffer containing 20 mM Tris pH 7.4, 140 mM NaCl, and 5% glycerol. Control enzymes with well-characterised cleavage activities, including Nth and Fpg **(New England Biolabs)**, were used for comparison. Reactions initiated mixing the identical volumes of 2-fold concentrated premixes to achieve 40nM DNA substrate and 70 nM enzyme (unless otherwise stated). Reactions were quenched by the addition of formamide at specific time points. Reaction products were separated on 15% denaturing polyacrylamide gels (containing 15% acrylamide/bisacrylamide [19:1], 8 M urea, and TBE buffer). Assays included mNeil3 reactions with both its Ap-ICL substrate and DNA oligonucleotides containing Ap sites.

### Trapping assay

Experiments, including trapping DNA with Na[BH₃(CN)] were performed in the same buffer as described above. Firstly, DNA and mNeil3 were mixed and incubated at room temperature. Secondly, Na[BH₃(CN)] was added to trap reaction intermediates. This reaction was terminated after 5 minutes by the addition of formamide, and separated on 15% denaturing gel. Activity measurements of the M1, V2M, and K82A mNeil3 variants were performed in a reaction buffer containing 20mM Tris pH 7.4, 140mM NaCl, 5% glycerol and 1mM TCEP. Four distinct DNA substrates were used in assays with the M1 and V2M enzymes, with the specific substrate combinations described in the Supplementary Figure (Figure 1 + Supplementary Figure 2). Proteins were mixed with the best selected DNA substrate Ap-ICL, and reactions were stopped at several time points (Figure 2). Also, protein concentrations were varied (Supplementary Figure 1). Stopped reactions were separated on both a 15 % denaturing acrylamide gel and SDS gel.

**Figure 1:**
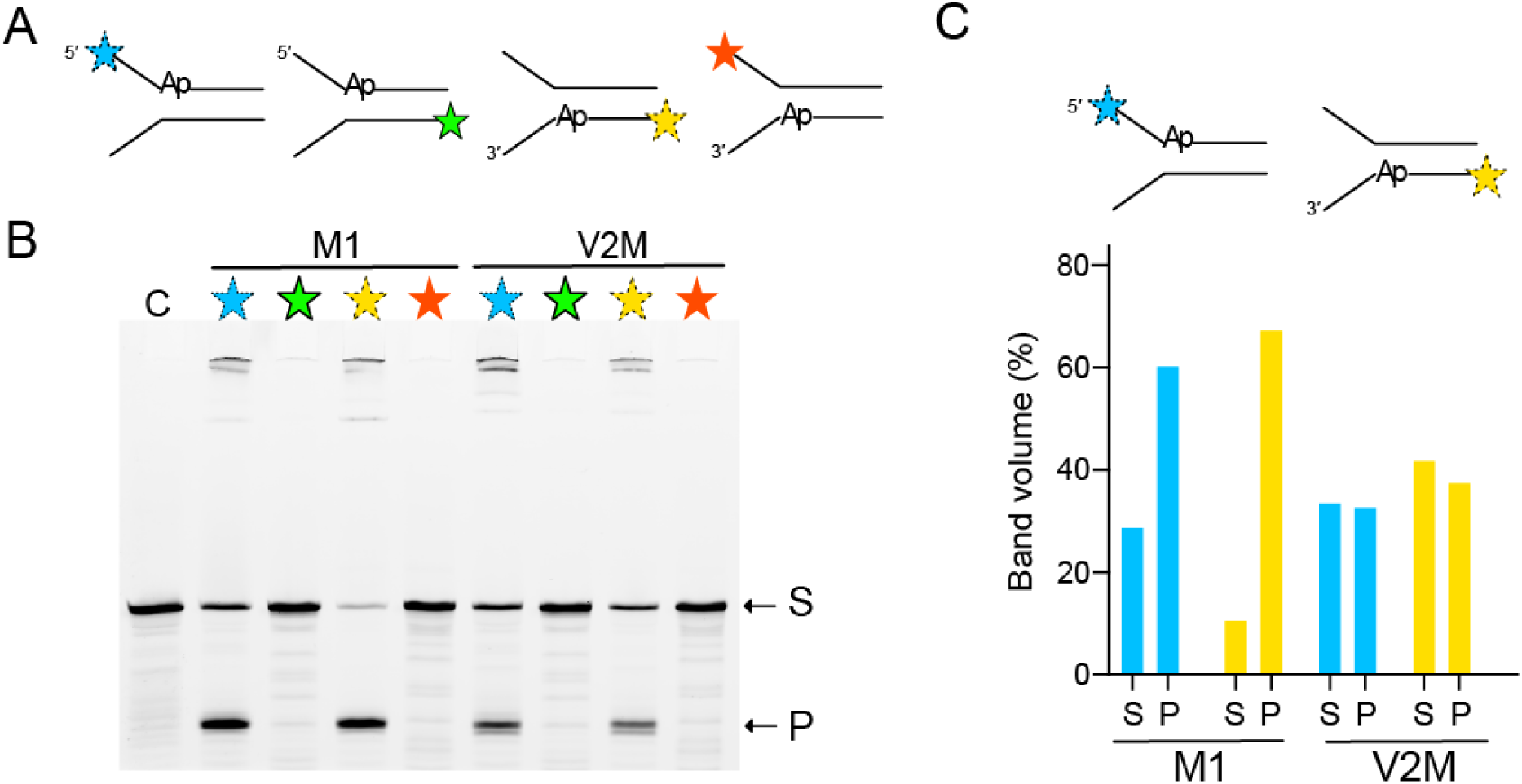
The V2M NEI domain has a weaker lyase activity toward the Ap site compared to the M1 NEI (A) Schematic representation of DNA substrates containing an Ap site positioned at different locations. The star indicates the position of the ATTO488 fluorescent label, with distinct star colours used to distinguish among the individual substrates. (B) 15% denaturing PAGE gel of enzymatic reactions of DNA substrates containing Ap site and Neil3 with either M1 or V2M on the first position. Stars correspond to the appropriate DNA substrate. (C) Band volumes of substrate S and product P for two DNA substrates were depicted in a bar chart.

**Figure 2:**
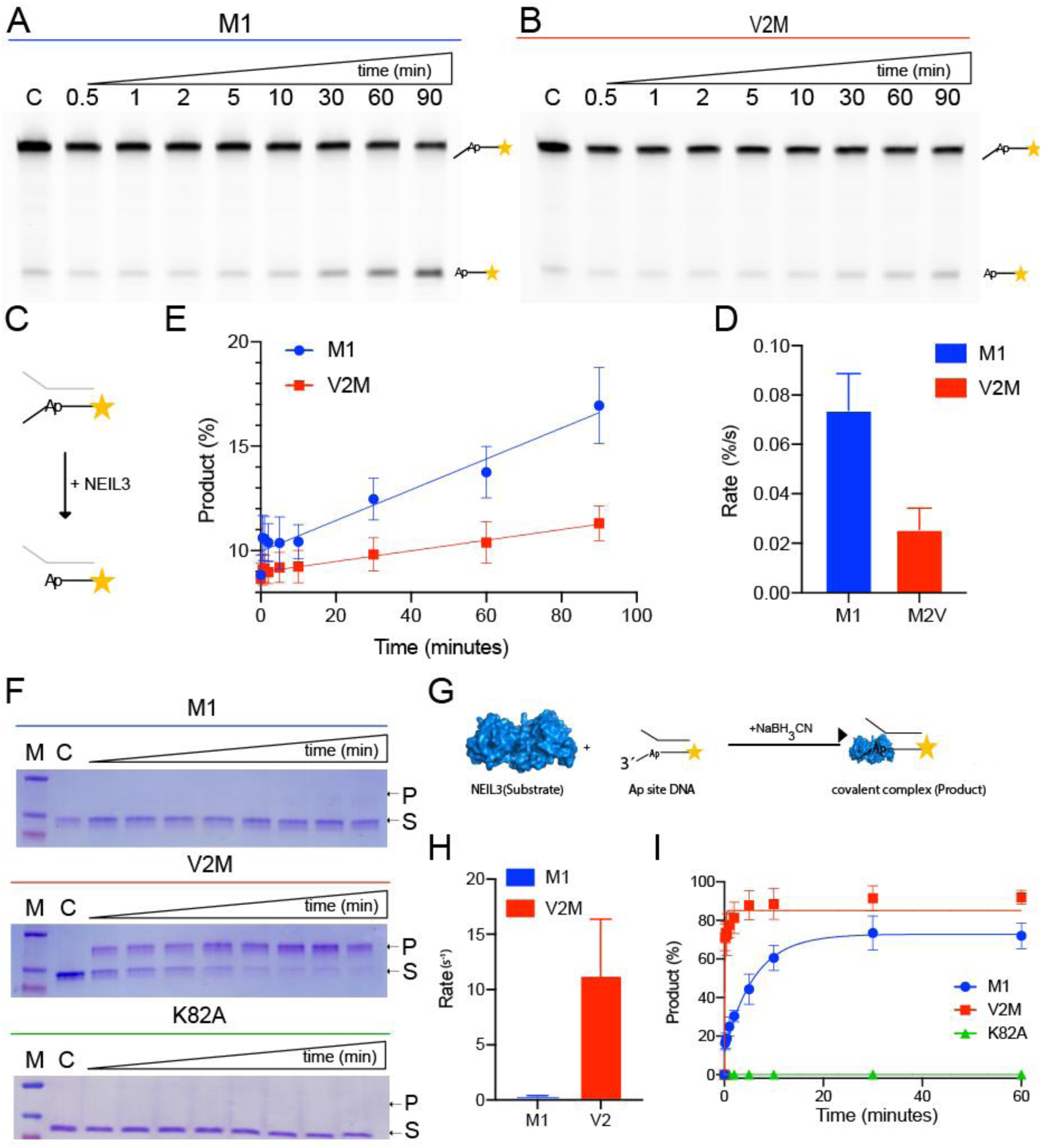
Processing of an Ap-site by Neil3 glycosylase variants M1 NEI and V2M NEI. Time-course comparison of Neil3 variant M1 NEI (A) and V2M NEI (B) with DNA substrate containing Ap-site. (C) Reaction scheme of the enzymatic assay. Gels were analysed, and product percentages were plotted in a graph (D), and the rate was calculated and depicted in a bar chart (E). Neil3 variant M1 NEI, V2M NEI and mutant K82A were tested in trapping assays with illustrative gels shown in (F) with the reaction scheme depicted in (G). The percentage of band volume corresponding to the covalent product was depicted in a graph for each Neil3 variant (H), and the rate was calculated and depicted in a bar chart (I).

### Anisotropy binding assay

ATTO488 labelled oligonucleotide containing THF (tetrahydrofuran) analogue that mimics Ap site was custom synthesised (Eurofins, sequence is listed in Supplementary Figure 2). For binding assays oligonucleotide was resuspended in a reaction buffer composed of 20 mM Tris pH = 7.4, 140 mM NaCl, 0.5 mM TCEP and 5% glycerol. DNA substrate was titrated with pure mNeil3 M1 and K82A from 0 µM to 2 uM, separated on 15% native PAGE and visualised on TYPHOON FLA 7000 fluorescence imager (GE Healthcare). Gels were qualitative control of DNA binding, completing fluorescence anisotropy. No loading dye was used for native PAGE, only 10% glycerol.

To determine K_D_, we used the fluorescence anisotropy measurements in an identical reaction buffer and with the same substrates as described in gel-based assays. The concentration of ATTO488-labelled DNA was 50 nM in the reaction buffer. This reaction mixture was titrated with mNeil3, and K_D_ was calculated as described before (24).

## Results

### V2M NEI domain exhibits weaker Ap lyase activity compared to M1 NEI

We assessed the Ap lyase activity of both variants using a panel of fork substrates. As expected, the non-Ap strand (labelled but lesion-free) showed no cleavage. M1 NEI displayed higher Ap lyase activity than V2M NEI (Figure 1B), cleaving more than 60% of the substrate, whereas V2M NEI cleaved only ∼40%. M1 NEI preferentially cleaved substrates in which the Ap site was positioned on a fork bearing a single-stranded 3′ end. In contrast, V2M NEI showed more uniform behaviour as the activity across substrates containing Ap sites at distinct positions was similar (Figure 1C). These findings prompted a direct comparison of the kinetic and mechanistic properties of the M1 and V2M forms, particularly in their processing of Ap sites versus Ap-ICLs.

### V2M NEI forms a covalent intermediate rapidly and avidly but cleaves Ap sites more reluctantly than M1 NEI

We next compared the enzyme kinetics of the two NEI variants using a 3′-Ap fork substrate (Figure 2C). Time-course analysis (Figure 2A–E) revealed that M1 NEI cleaved Ap sites approximately threefold faster than V2M NEI (Figure 2E), as determined by quantifying product accumulation over time (Figure 2F). To investigate whether this discrepancy arose from differences in catalytic turnover or substrate binding, we performed Na[BH₃(CN)] trapping assays, which stabilise the covalent enzyme–DNA intermediate formed during nucleophilic attack (Figure 2H). V2M NEI consistently formed a greater amount of trapped covalent complex under identical conditions than M1 NEI (Figure 2G, I), indicating that V2M engages with Ap site and forms the covalent intermediate more readily, as a result of faster initial binding to the Ap site. As expected, the catalytically inactive K82A mutant failed to form trapped complexes. Together, these data indicate that V2M NEI exhibits efficient lesion engagement and intermediate formation but progresses more slowly through the lyase reaction. This results in reduced overall turnover and lower Ap-lyase efficiency compared with M1 NEI.

### V2M NEI preferentially unhooks Ap-ICLs without strand cleavage

To examine how these NEI variants process the biologically relevant abasic interstrand crosslink (Ap-ICL), we compared their activity using an Ap-ICL-containing fork substrate. Titration experiments demonstrated that V2M NEI processed Ap-ICLs more efficiently than M1 NEI at all tested concentrations, achieving complete substrate conversion at 560 nM (Supplementary Figure 1). Time-course analysis further revealed qualitative differences in product formation. V2M NEI predominantly generated product 1, corresponding to unhooked DNA lacking a single-strand break (Figure 3A–E). In contrast, M1 NEI favoured formation of product 2, a lyase-dependent β-elimination product associated with strand cleavage, and processed Ap-ICLs more slowly overall. These data indicate that M1 NEI preferentially channels Ap-ICL processing into strand-cleaving reactions, whereas V2M NEI favours glycosylase-mediated unhooking without introducing breaks.

**Figure 3:**
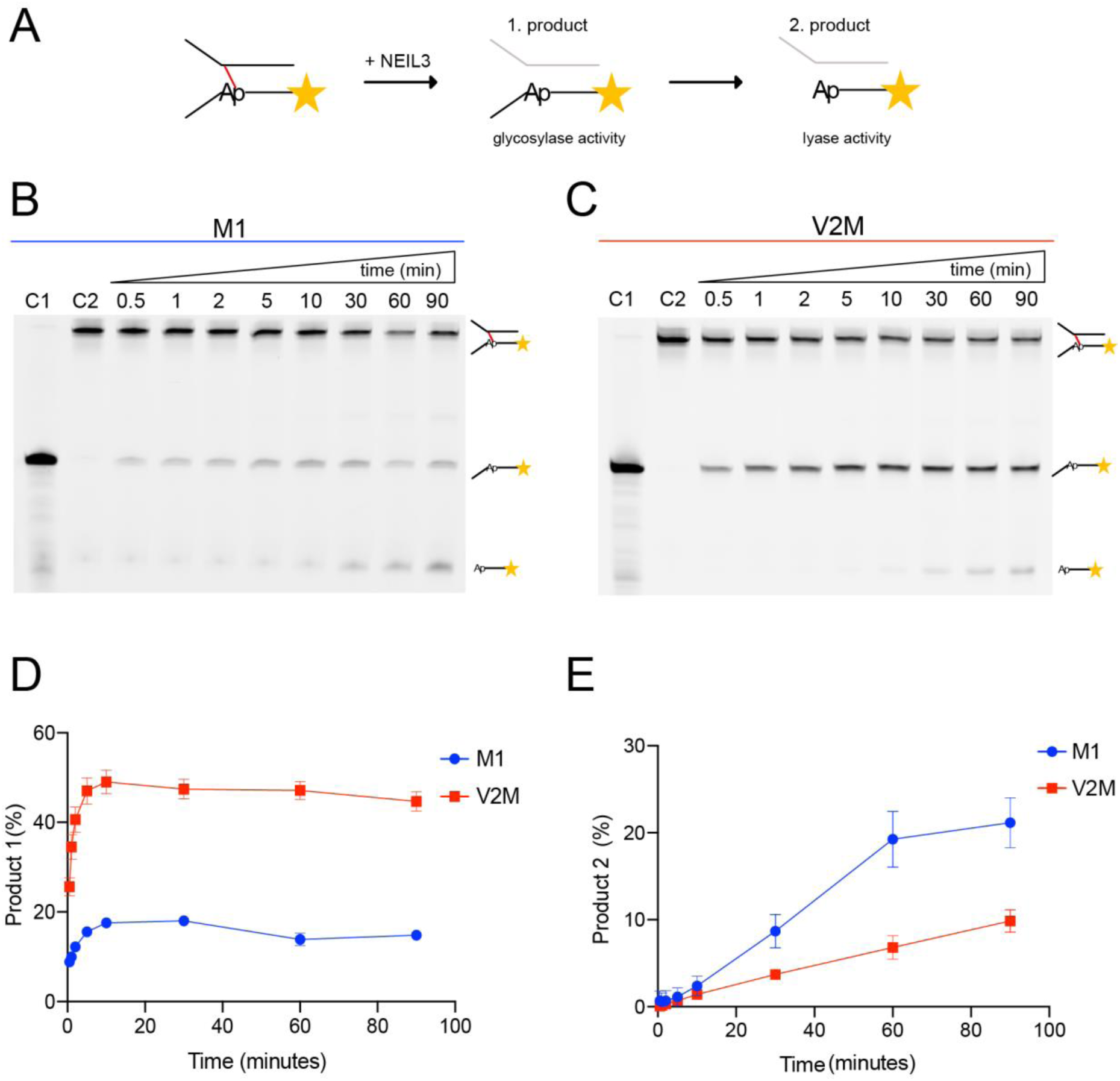
Processing of Ap-ICL by Neil3 glycosylase variants M1 and V2M NEI. (A) Scheme of Neil3 glycosylase and lyase activity on Ap-ICL and the formed products. Glycosylase and lyase activity of Neil3 variants M1 (B) and V2M (C) Nei were tested in an enzymatic assay with a DNA fork containing Ap-ICL. Each time point was resolved on the 15% denaturing PAGE gel. Gels were analysed, and product percentage from glycosylase (D, product 1) and lyase (E, product 2) activity was plotted in a graph.

### Neil3 lyase products of Ap-ICL correspond to β,δ-elimination chemistry

Members of the Fpg/Nei family possess both DNA glycosylase and Ap lyase activities, with β,δ-elimination generating strand breaks bearing 3′-phosphate termini. To define the chemical nature of Neil3 cleavage products, we incubated Ap-site and Ap-ICL substrates with Neil3 and analysed reaction products by denaturing PAGE (Figure 4A). Identical substrate sequences were used throughout. Control reactions with Fpg and Nth confirmed generation of 3′-phosphate and 3′-dRP termini (deoxyribose phosphate), respectively (37, 38). All Neil3 reactions produced products co-migrating with 3′-phosphate, consistent with β-elimination chemistry analogous to Fpg. Progression to δ-elimination was inefficient under the conditions tested, indicating that while Neil3 is capable of canonical β,δ-lyase chemistry, strand-break formation remains limited, particularly during Ap-ICL processing by V2M NEI.

**Figure 4:**
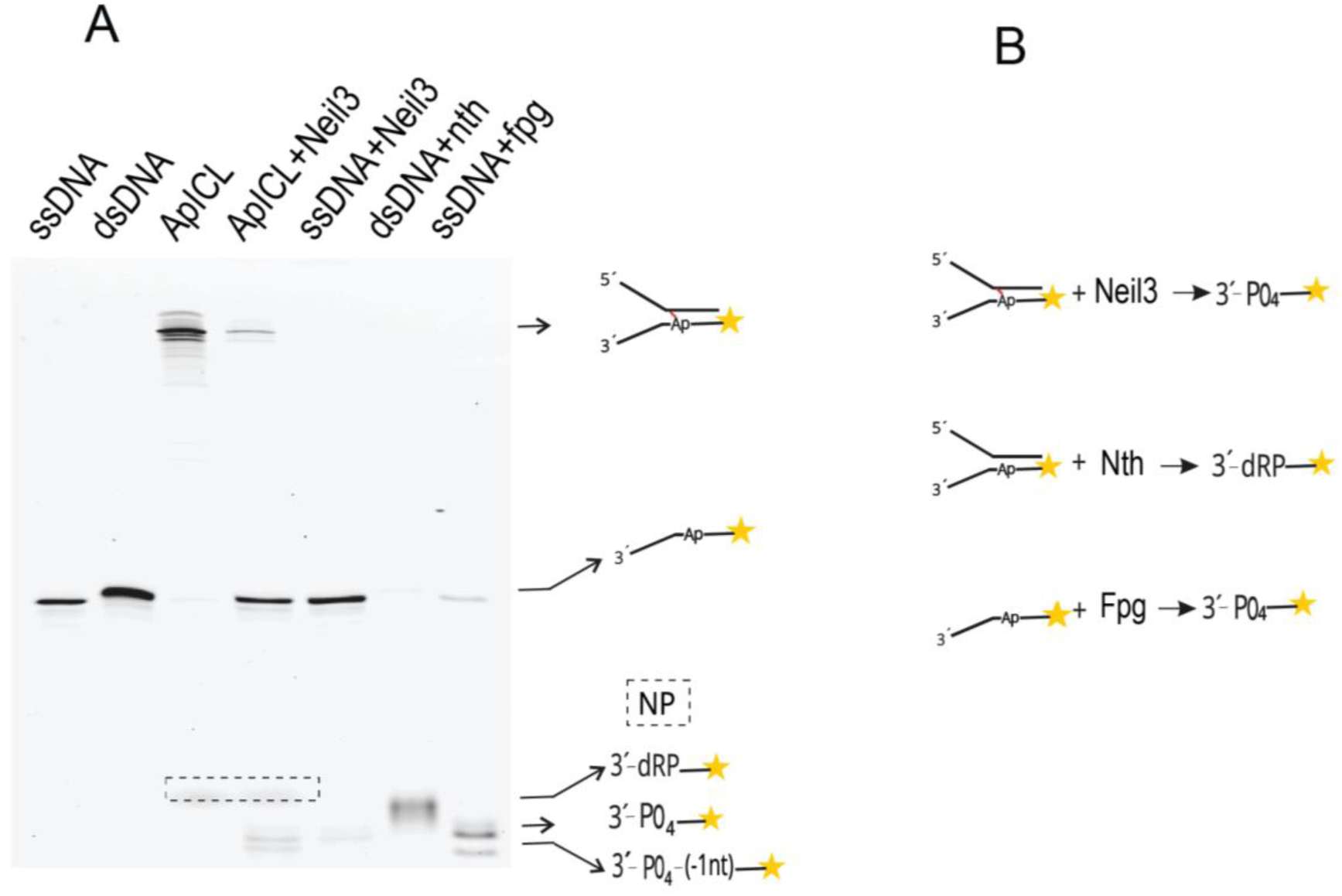
Enzymatic reaction reveals the product of Neil3 lyase activity. (A) The formamidopyrimidine-DNA glycosylase (Fpg) and Endonuclease (Nth) were used to compare final cleavage products of Neil3. Reactions with different substrates were separated on 15% denaturing PAGE. For clarity, products in the box were saturated and stretched for easier comparison. (B) For both those enzymes, the substrate and the product of their cleavage are known.

### Ap-ICL and Ap site repair intermediates upon timed crosslinker addition

Given the similar chemical nature of Ap-site and Ap-ICL products, we next examined whether Neil3 processes these substrates through comparable covalent intermediates. Using Na[BH₃(CN)] trapping, we analysed reaction intermediates formed during Neil3 activity on both substrates (Figure 5). When Na[BH₃(CN)] was added at the start of the reaction, Ap-site substrates yielded the expected stable covalent intermediate (Figure 5, lane 10). In contrast, Ap-ICL reactions were arrested without formation of a trapped intermediate under identical conditions (Figure 5, lane 4). To resolve this discrepancy, we performed time-resolved trapping experiments in which Na[BH₃(CN)] was added at defined time points after reaction initiation (Figure 5, lanes 5–9). Under these conditions, enzyme–DNA adducts became detectable, including a major species co-migrating with the Ap-site intermediate and additional higher-molecular-weight transient species observed specifically during Ap-ICL processing (Figure 5, red box). The size and migration of the dominant trapped intermediate matched that of the crystallised NEI–Ap DNA complex.

**Figure 5:**
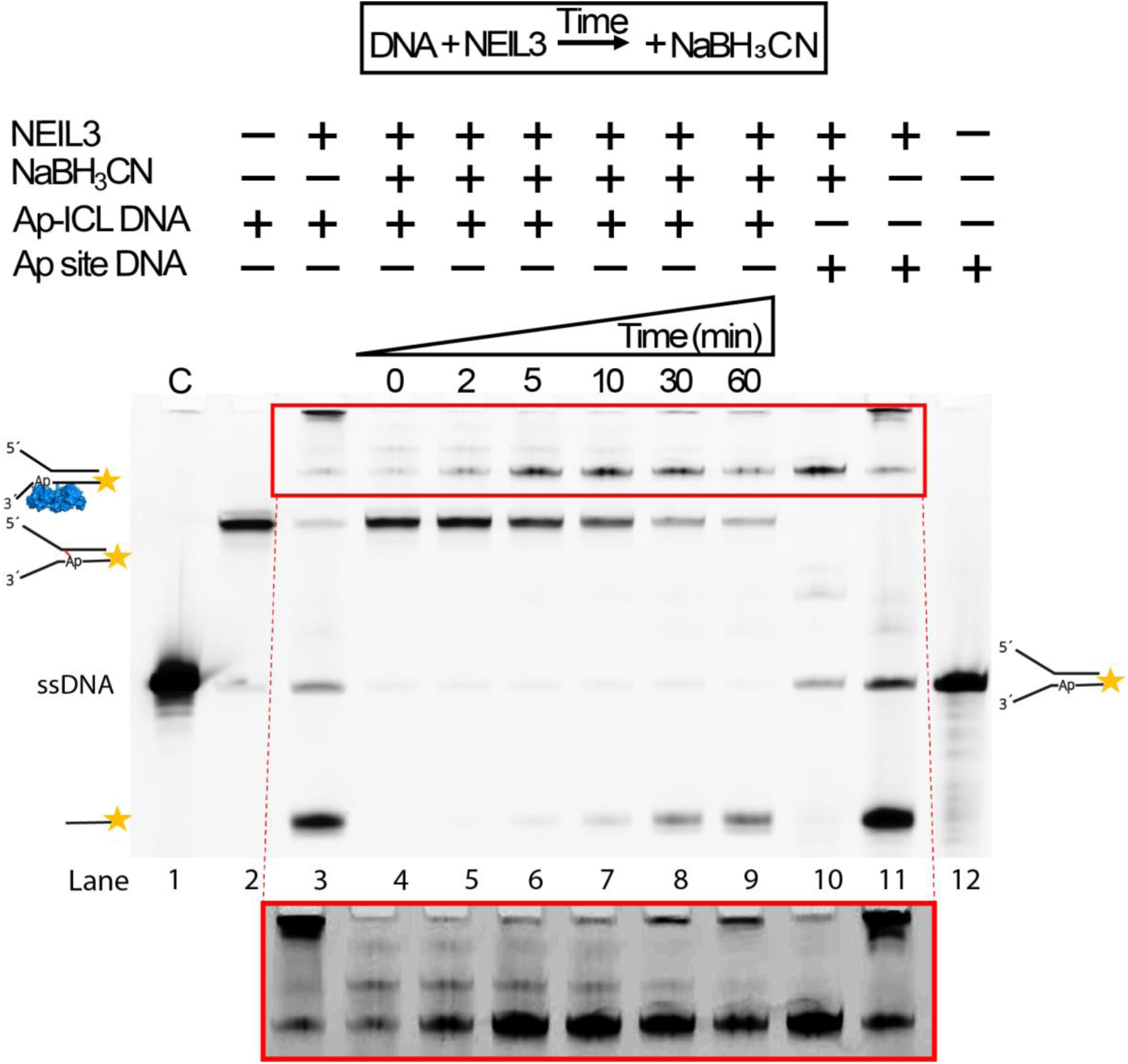
Time-resolved Na[BH₃(CN)] addition during Ap-ICL processing reveals enzyme–DNA complexes. The reaction of NEI domain mediated Ap-ICL repair with fluorescently labelled Ap-ICL on 5′ of Ap site containing DNA strand was subjected to Na[BH₃(CN)] trapping its reaction intermediates. Ap site reaction is shown as a control and for comparison. Na[BH₃(CN)] was either added at the start of the reaction or at a given time point of the time-course titration. Covalent adducts and higher migrating bands are highlighted within a red box.

### Crystal structure of NEI domain bound to ssDNA

We crystallised the mouse NEIL3 NEI domain (residues 1–282) covalently linked to an Ap site–containing single-stranded DNA substrate (PDB 8B9N). The Schiff-base linkage between the N-terminal amino group and the abasic sugar was stabilised by Na[BH₃(CN)] reduction prior to crystallisation. The overall fold of the NEI domain closely matches previously reported NEIL3 structures obtained with DNA hairpins or duplex-derived substrates (e.g., PDB 7Z5A, 3W0F), with an RMSD of 0.292 Å over non-hydrogen atoms (26, 35). In contrast to earlier trapped complexes, which represent unfavourable DNA structure, our structure captures NEIL3 engaged with a native single-stranded DNA substrate consistent with its replication-associated activity. Notably, we were able to observe the orientation of the bases in the ssDNA. Two nucleotides at the 3′ end showed well-defined electron density, these bases were rotated by approximately 90° relative to their orientation in duplex DNA eg. in previously reported NEI–DNA complexes. This unusual positioning appears to be stabilised by a cation-π pair stacking interaction with Arg262 (Figure 6). The observed DNA geometry supports a model in which NEIL3 engages lesions presented in a single-stranded context of replication forks.

**Figure 6:**
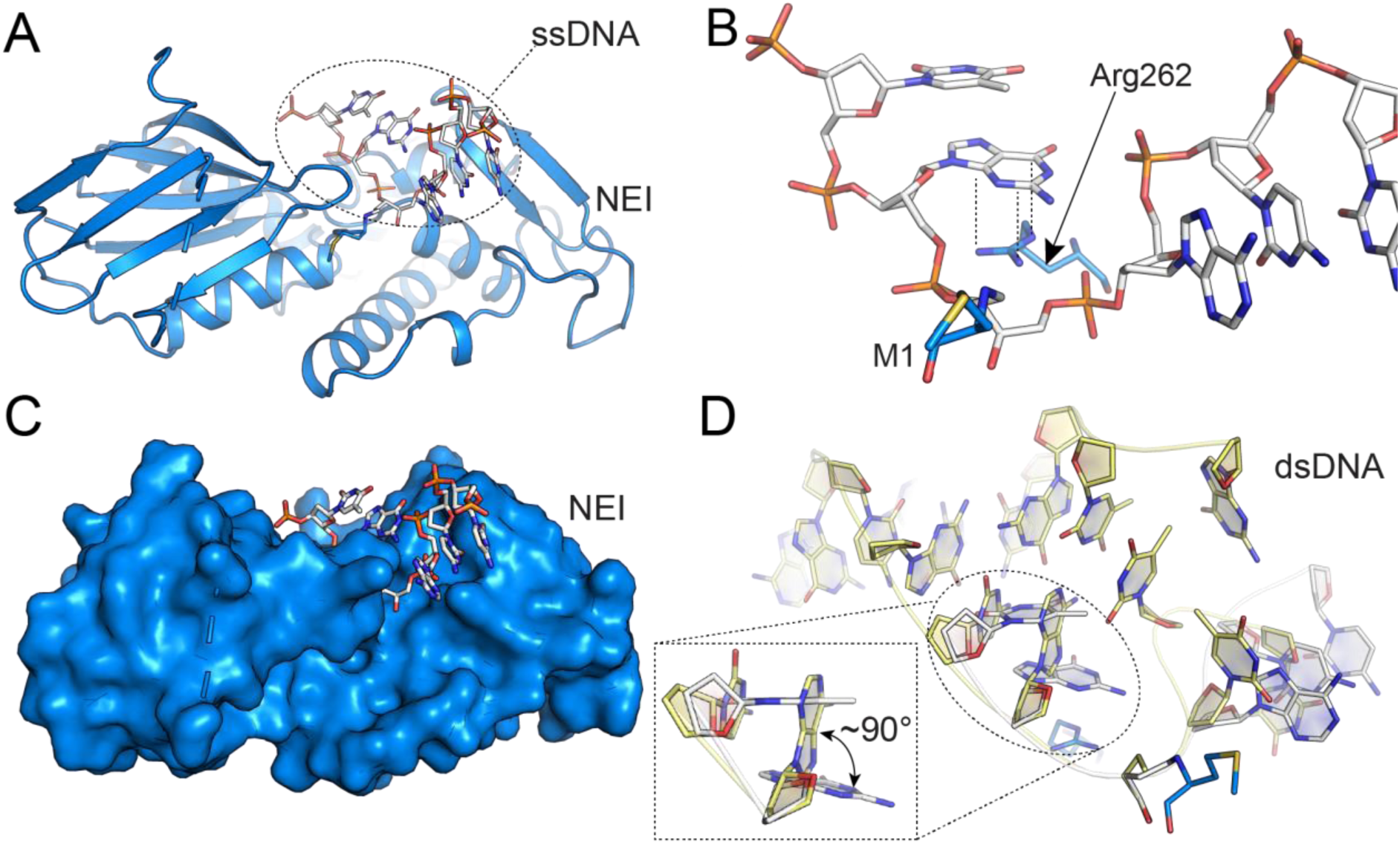
NEI structure with single-stranded DNA (covalently trapped reaction intermediate). (A) Overall structure of mouse NEIL3 NEI domain (M1) with ssDNA covalently trapped via Ap-site. (B) detail of M1 - Ap site linkage and π-ion interaction between DNA and Arg262 (C) Surface representation of NEI domain with DNA in active site. (D) An overlay of double-stranded DNA hairpin bound to NEI (PDB 7Z5A), conformational changes induced in ssDNA by binding to the glycosylase domain encircled by dotted line. DNA nucleobases on the 3′ site of the Ap site undergo rotation by ∼90° due to the interaction with Arg262 (PDB 8B9N).

## Binding of the K82A catalytic mutant to ssDNA

To assess whether loss of catalytic activity in the K82A mutant reflects impaired DNA binding, we measured substrate affinity using fluorescence anisotropy and electrophoretic mobility shift assays (EMSA) with a tetrahydrofuran (THF) abasic-site analogue, which preserves the DNA backbone architecture while lacking the reactive C1′ hydroxyl group. Despite complete loss of catalytic activity, K82A retained binding affinity comparable to wild-type mNEIL3 M1 (Figure 7B), indicating that K82 is required for catalysis rather than substrate engagement. This is consistent with K82 interacting primarily with the phosphodiester backbone rather than the abasic sugar itself, such that DNA binding remains representative despite the absence of the C1′ hydroxyl. EMSA confirmed efficient ssDNA binding by both wild-type and mutant proteins (Figure 7C).

**Figure 7:**
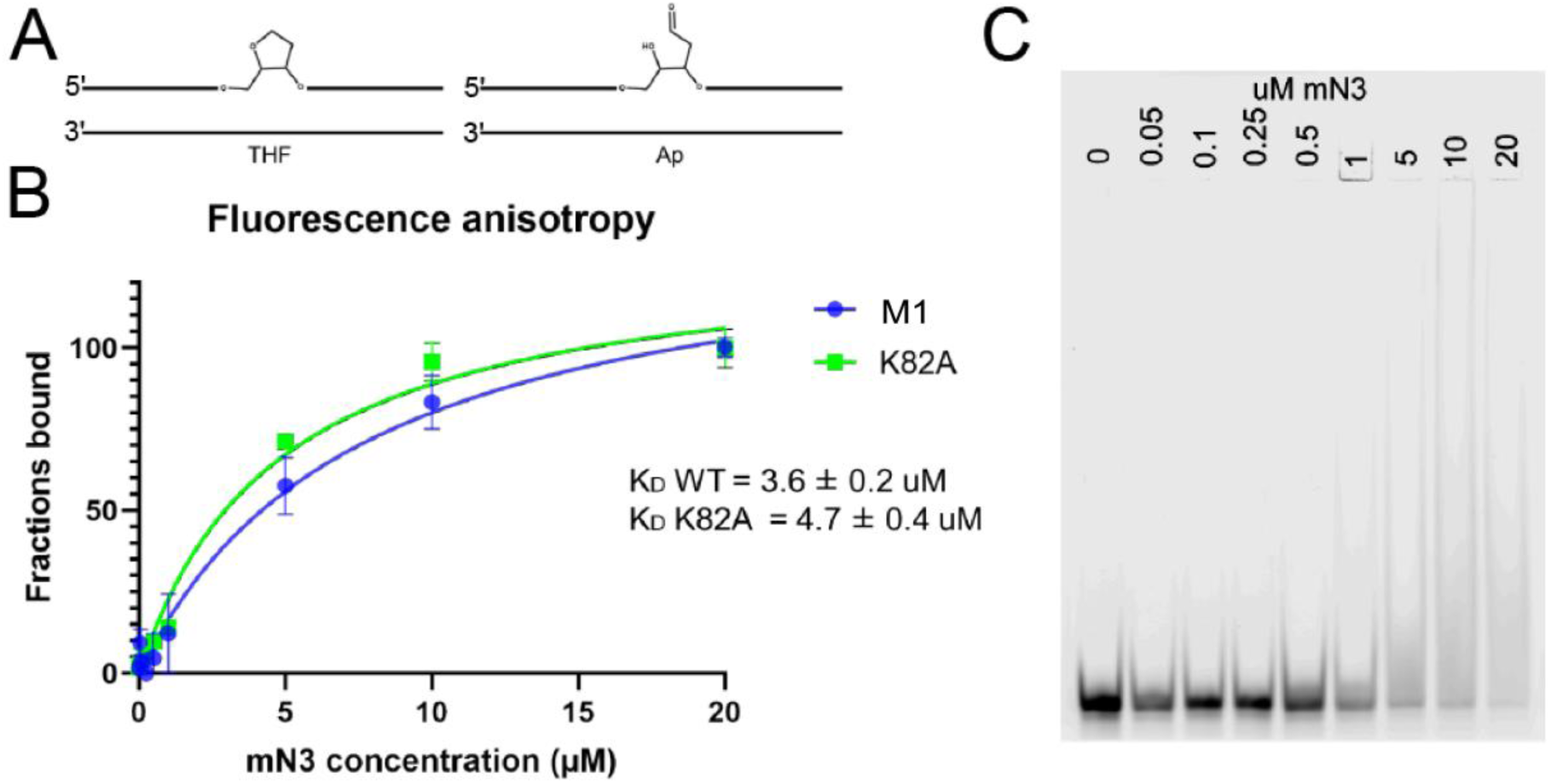
Fluorescence anisotropy binding assay. (A) Schematic representation of substrates containing the Ap site mimicking THF (tetrahydrofuran). (B) Comparison of affinities of mouse NEI M1 and NEI K82A mutant to single-stranded DNA with K_D_ values. (C) Electrophoretic mobility shift assay shows semiquantitative representation of interaction between ssDNA and NEI M1.

## Discussion

NEIL3 functions as a specialised replication-coupled enzyme that resolves abasic site interstrand crosslinks (Ap-ICLs) at stalled replication forks without generating double-strand breaks (DSBs). Its recruitment is triggered by TRAIP-dependent ubiquitylation of the CMG helicase, positioning NEIL3 precisely at sites of fork convergence during S phase (17, 19). *Xenopus* egg extract studies indicate that converging forks generate an X-shaped DNA structure that presents the Ap-ICL in a predominantly single-stranded context, providing a structural explanation for NEIL3’s strong preference for forked and ssDNA substrates (17, 35, 39).When NEIL3 activity is compromised, Ap-ICLs are rerouted to the Fanconi anaemia (FA) pathway, which relies on nucleolytic incision and DSB formation, underscoring that FA-mediated repair functions primarily as a backup rather than a redundant pathway(39, 40).

Our data provide a biochemical framework that explains how NEIL3 can uncouple Ap-ICLs without inducing strand breaks. The NEI catalytic domain mediates lesion engagement and unhooking, while auxiliary zinc finger and GRF domains stabilise ssDNA at the fork junction and orient the enzyme correctly relative to the lesion (26, 35). This division of individual labour likely prevents steric competition between DNA binding and catalysis at ssDNA-rich replication intermediates and distinguishes NEIL3 from canonical Fpg/Nei glycosylases that act primarily on duplex DNA(6). Structural evidence supporting ssDNA-specific engagement reinforces the view that NEIL3 has evolved away from global base excision repair toward replication-associated lesion processing (10).

A central finding of this study is that the catalytic outcome of NEIL3 is dictated by the N-terminal processing state of the NEI domain. The V2M variant, which mimics native V2, efficiently unhooks Ap-ICLs while strongly attenuating Ap-lyase–mediated strand cleavage. In contrast, the unprocessed M1 variant favours lyase activity and generates ssDNA breaks. This resolves the longstanding discrepancy between biochemical reports of NEIL3 lyase activity forming a strand break in the Ap-site DNA strand and studies from JC Walter laboratory showing incision-free repair(10, 27, 36). Our results support earlier implicit proposals that V2 constitutes the catalytically relevant N-terminal residue and demonstrate that incomplete N-terminal processing during recombinant expression can profoundly bias observed enzyme activity.

Time-resolved Na[BH₃(CN)] trapping has revealed that Ap-ICL unhooking by NEIL3 proceeds through additional transient reaction intermediates that are distinct from those formed during processing of isolated Ap sites. Unlike Ap-site substrates, which yield a single, stable trapped (reduced) Schiff-base intermediate characteristic of Fpg/Nei enzymes, Ap-ICL substrates produce short-lived, higher-molecular-weight covalent species and largely prevent progression to the canonical trapped product, the Schiff-base intermediate, which appears only at later stages of the reaction (41–43). The appearance, migration, and limited lifetime of these species suggest a more complex reaction mechanism that either precedes or bypasses formation of the classical Schiff-base intermediate. This behaviour provides a mechanistic snapshot of NEIL3 specialisation for interstrand crosslink repair, enabling lesion unhooking while preserving replication fork integrity, and helps reconcile *in vitro* biochemical data with replication-coupled repair models derived from extract systems. We outline the proposed reaction mechanism for Ap-ICL repair by NEIL3 glycosylase in Figure 8.

**Figure 8:**
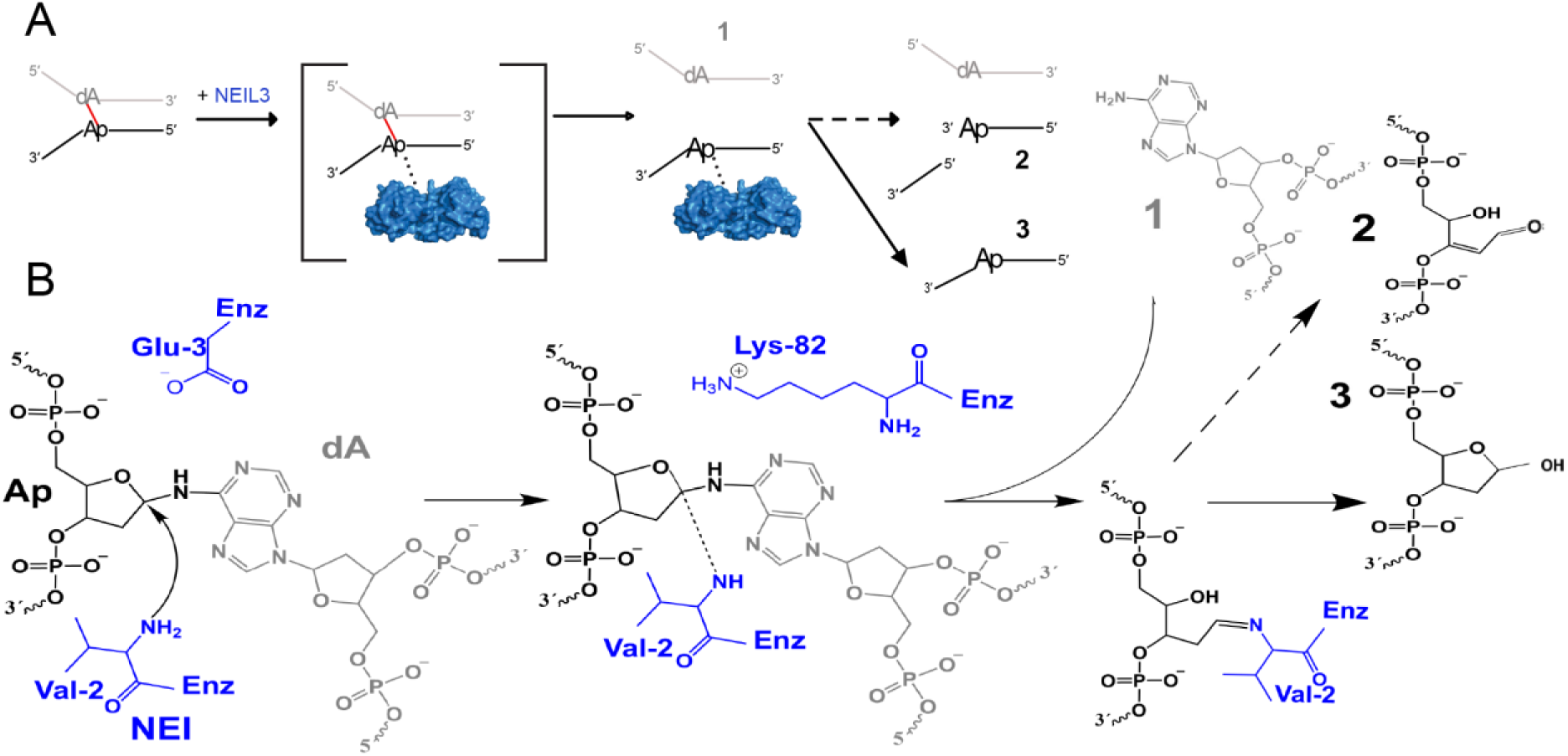
Mechanism of Neil3 Ap-ICL repair. (A) Illustrative scheme of Neil3 Ap-ICL repair. (B) The N-terminal amino group attacks the non-native N-glycosidic bond of the AP-ICL, with this nucleophilic attack stabilized by Glu-3. Unhooking of the AP-ICL appears to depend on Lys-82, which is directly involved in catalysis, as mutation to alanine abolishes enzymatic activity while DNA binding is largely maintained (Figure 7). The opposing adenine of the crosslink is released as undamaged ssDNA (product 1). Dissociation of the enzyme generates product 3, single-stranded AP site DNA, which is the preferred product of V2M NEI, with the AP sugar mostly in the hemiacetal (closed-ring) form. Alternatively, undesired lyase activity produces product 2, a DNA strand break with a reactive 3′-dRP (open-chain aldehyde) at the 3′ terminus, which predominates for M1 NEI.

Together, these findings establish NEIL3 as a highly specialised glycosylase whose activity is tuned to replication-associated DNA structures. By coupling efficient Ap-ICL unhooking with suppression of strand cleavage, NEIL3 enables lesion bypass and replication restart while minimising the risk of DSB formation, thereby defining a distinct, non-nucleolytic pathway for endogenous crosslink repair.

## DATA AVAILABILITY

The structural data generated for this study have been deposited into the Protein Data Bank under the PDB code 8B9N. All data analyses are presented in the main text or supplementary materials and supplementary data. All source data underlying the graphs and charts presented in the main figures are presented as Supplementary Data (either as standalone graphs, excel tables or in text format).

## SUPPLEMENTARY DATA

Supplementary Data are available at NAR Online.

## REFERENCES

1. Lindahl, T. (1993) Instability and decay of the primary structure of DNA. Nature, 362, 709–715.

2. Lindahl, T. and Andersson, a (1972) Rate of chain breakage at apurinic sites in double-stranded deoxyribonucleic acid. Biochemistry, 11, 3618–3623.

3. Lindahl, T. (1974) An N-Glycosidase from Escherichia coli That Releases Free Uracil from DNA Containing Deaminated Cytosine Residues. Proceedings of the National Academy of Sciences, 71, 3649–3653.

4. Roos, W.P. and Kaina, B. (2013) DNA damage-induced cell death: From specific DNA lesions to the DNA damage response and apoptosis. Cancer Lett., 332, 237–248.

5. Fromme, J.C. and Verdine, G.L. (2004) Base excision repair. Adv. Protein Chem., 69, 1–41.

6. Nagorska, K., Silhan, J., Li, Y., Pelicic, V., Freemont, P.S., Baldwin, G.S. and Tang, C.M. (2012) A network of enzymes involved in repair of oxidative DNA damage in Neisseria meningitidis. Mol. Microbiol., 83, 1064–1079.

7. Price, N.E., Johnson, K.M., Wang, J., Fekry, M.I., Wang, Y. and Gates, K.S. (2014) Interstrand DNA–DNA Cross-Link Formation Between Adenine Residues and Abasic Sites in Duplex DNA. J. Am. Chem. Soc., 136, 3483–3490.

8. Johnson, K.M., Price, N.E., Wang, J., Fekry, M.I., Dutta, S., Seiner, D.R., Wang, Y. and Gates, K.S. (2013) On the formation and properties of interstrand DNA-DNA cross-links forged by reaction of an abasic site with the opposing guanine residue of 5′-CAp sequences in duplex DNA. J. Am. Chem. Soc., 135, 1015–1025.

9. Housh, K., Jha, J.S., Yang, Z., Haldar, T., Johnson, K.M., Yin, J., Wang, Y. and Gates, K.S. (2021) Formation and Repair of an Interstrand DNA Cross-Link Arising from a Common Endogenous Lesion. J. Am. Chem. Soc., 143, 15344–15357.

10. Imani Nejad, M., Housh, K., Rodriguez, A.A., Haldar, T., Kathe, S., Wallace, S.S., Eichman, B.F. and Gates, K.S. (2020) Unhooking of an interstrand cross-link at DNA fork structures by the DNA glycosylase NEIL3. DNA Repair (Amst)., 86, 102752.

11. Huskova, A., Landova, B., Boura, E. and Silhan, J. (2022) The rate of formation and stability of abasic site interstrand crosslinks in the DNA duplex. DNA Repair (Amst)., 113, 103300.

12. Noll, D.M., McGregor Mason, T., Miller, P.S., Mason, T.M. and Miller, P.S. (2006) Formation and repair of interstrand cross-links in DNA. Chem Rev, 106, 277–301.

13. Roos, W.P. and Kaina, B. (2006) DNA damage-induced cell death by apoptosis. Trends Mol. Med., 12, 440–450.

14. Karmakar, S., Purkait, K., Chatterjee, S. and Mukherjee, A. (2016) Anticancer activity of a cis-dichloridoplatinum(II) complex of a chelating nitrogen mustard: insight into unusual guanine binding mode and low deactivation by glutathione. Dalton Transactions, 45, 3599–3615.

15. Räschle, M., Knipsheer, P., Enoiu, M., Angelov, T., Sun, J., Griffith, J.D., Ellenberger, T.E., Schärer, O.D. and Walter, J.C. (2008) Mechanism of Replication-Coupled DNA Interstrand Crosslink Repair. Cell, 134, 969–980.

16. Knipscheer, P., Räschle, M., Smogorzewska, A., Enoiu, M., Ho, T.V., Schärer, O.D., Elledge, S.J. and Walter, J.C. (2009) The fanconi anemia pathway promotes replication-dependent DNA interstrand cross-link repair. Science (1979)., 326, 1698–1701.

17. Semlow, D.R., Zhang, J., Budzowska, M., Drohat, A.C. and Walter, J.C. (2016) Replication-Dependent Unhooking of DNA Interstrand Cross-Links by the NEIL3 Glycosylase. Cell, 167, 498–511.e14.

18. Hodskinson, M.R.G., Bolner, A., Sato, K., Kamimae-Lanning, A.N., Rooijers, K., Witte, M., Mahesh, M., Silhan, J., Petek, M., Williams, D.M., et al. (2020) Alcohol-derived DNA crosslinks are repaired by two distinct mechanisms. Nature, 579, 603–608.

19. Wu, R.A., Semlow, D.R., Kamimae-Lanning, A.N., Kochenova, O. V, Chistol, G., Hodskinson, M.R., Amunugama, R., Sparks, J.L., Wang, M., Deng, L., et al. (2019) TRAIP is a master regulator of DNA interstrand crosslink repair. Nature, 567, 267–272.

20. Zharkov, D.O., Shoham, G. and Grollman, A.P. (2003) Structural characterization of the Fpg family of DNA glycosylases. DNA Repair (Amst)., 2, 839–862.

21. Bhagwat, M. and Gerlt, J. a (1996) 3’- and 5’-strand cleavage reactions catalyzed by the Fpg protein from Escherichia coli occur via successive beta- and delta-elimination mechanisms, respectively. Biochemistry, 35, 659–665.

22. Coste, F., Ober, M., Carell, T., Boiteux, S., Zelwer, C. and Castaing, B. (2004) Structural basis for the recognition of the FapydG lesion (2,6-diamino-4-hydroxy-5-formamidopyrimidine) by formamidopyrimidine-DNA glycosylase. Journal of Biological Chemistry, 279, 44074–44083.

23. Fromme, J.C. and Verdine, G.L. (2002) Structural insights into lesion recognition and repair by the bacterial 8-oxoguanine DNA glycosylase MutM. Nat. Struct. Biol., 9, 544–552.

24. Landova, B. and Silhan, J. (2020) Conformational changes of DNA repair glycosylase MutM triggered by DNA binding. FEBS Lett., 594, 3032–3044.

25. Jiang, D., Hatahet, Z., Melamede, R.J., Kow, Y.W. and Wallace, S.S. (1997) Characterization of Escherichia coli Endonuclease VIII*. Journal of Biological Chemistry, 272, 32230–32239.

26. Liu, M., Imamura, K., Averill, A.M., Wallace, S.S. and Doublié, S. (2013) Structural characterization of a mouse ortholog of human NEIL3 with a marked preference for single-stranded DNA. Structure, 21, 247–256.

27. Yang, Z., Nejad, M.I., Varela, J.G., Price, N.E., Wang, Y. and Gates, K.S. (2017) A role for the base excision repair enzyme NEIL3 in replication-dependent repair of interstrand DNA cross-links derived from psoralen and abasic sites. DNA Repair (Amst)., 52, 1–11.

28. Takao, M., Oohata, Y., Kitadokoro, K., Kobayashi, K., Iwai, S., Yasui, A., Yonei, S. and Zhang, Q.M. (2009) Human Nei-like protein NEIL3 has AP lyase activity specific for single-stranded DNA and confers oxidative stress resistance in Escherichia coli mutant. Genes to Cells, 14, 261–270.

29. Liu, M., Bandaru, V., Bond, J.P., Jaruga, P., Zhao, X., Christov, P.P., Burrows, C.J., Rizzo, C.J., Dizdaroglu, M. and Wallace, S.S. (2010) The mouse ortholog of NEIL3 is a functional DNA glycosylase in vitro and in vivo. Proc. Natl. Acad. Sci. U. S. A., 107, 4925–4930.

30. Liu, M., Doublié, S. and Wallace, S.S. (2013) Neil3, the final frontier for the DNA glycosylases that recognize oxidative damage. Mutation Research - Fundamental and Molecular Mechanisms of Mutagenesis, 743–744, 4–11.

31. Liu, M., Bandaru, V., Holmes, A., Averill, A.M., Cannan, W. and Wallace, S.S. (2012) Expression and purification of active mouse and human NEIL3 proteins. Protein Expr. Purif., 84, 130–139.

32. Hirel, P.H., Schmitter, J.M., Dessen, P., Fayat, G. and Blanquet, S. (1989) Extent of N-terminal methionine excision from Escherichia coli proteins is governed by the side-chain length of the penultimate amino acid. Proc. Natl. Acad. Sci. U. S. A., 86, 8247–8251.

33. Frottin, F., Martinez, A., Peynot, P., Mitra, S., Holz, R.C., Giglione, C. and Meinnel, T. (2006) The proteomics of N-terminal methionine cleavage. Molecular and Cellular Proteomics, 5, 2336–2349.

34. Dalbøge, H., Bayne, S. and Pedersen, J. (1990) In vivo processing of N-terminal methionine in E. coli. FEBS Lett., 266, 1–3.

35. Huskova, A., Dinesh, D.C., Srb, P., Boura, E., Veverka, V. and Silhan, J. (2022) Model of abasic site DNA cross-link repair; from the architecture of NEIL3 DNA binding domains to the X-structure model. Nucleic Acids Res., 50, 10436–10448.

36. Rodriguez, A.A., Wojtaszek, J.L., Greer, B.H., Haldar, T., Gates, K.S., Williams, R.S. and Eichman, B.F. (2020) An autoinhibitory role for the GRF zinc finger domain of DNA glycosylase NEIL3. Journal of Biological Chemistry, 10.1074/jbc.ra120.015541.

37. Silhan, J., Nagorska, K., Zhao, Q., Jensen, K., Freemont, P.S., Tang, C.M. and Baldwin, G.S. (2012) Specialization of an Exonuclease III family enzyme in the repair of 3’ DNA lesions during base excision repair in the human pathogen Neisseria meningitidis. Nucleic Acids Res., 40, 2065–2075.

38. Silhan, J., Zhao, Q., Boura, E., Thomson, H., Förster, A., Tang, C.M., Freemont, P.S. and Baldwin, G.S. (2018) Structural basis for recognition and repair of the 3′-phosphate by NExo, a base excision DNA repair nuclease from Neisseria meningitidis. Nucleic Acids Res., 46, 11980–11989.

39. Semlow, D.R. and Walter, J.C. (2021) Mechanisms of Vertebrate DNA Interstrand Cross-Link Repair. Annu. Rev. Biochem., 90, 107–135.

40. Li, N., Wang, J., Wallace, S.S., Chen, J., Zhou, J. and D’Andrea, A.D. (2020) Cooperation of the NEIL3 and Fanconi anemia/BRCA pathways in interstrand crosslink repair. Nucleic Acids Res., 48, 3014–3028.

41. Tchout, J. and Grollman, A.P. (1995) The Catalytic Mechanism of Fpg Protein. Journal of Biological Chemistry, 270, 11671–11677.

42. Gilboa, R., Zharkov, D.O., Golan, G., Fernandes, A.S., Gerchman, S.E., Matz, E., Kycia, J.H., Grollman, A.P. and Shoham, G. (2002) Structure of formamidopyrimidine-DNA glycosylase covalently complexed to DNA. Journal of Biological Chemistry, 277, 19811–19816.

43. Sugahara, M., Mikawa, T., Kumasaka, T., Yamamoto, M., Kato, R., Fukuyama, K., Inoue, Y. and Kuramitsu, S. (2000) Crystal structure of a repair enzyme of oxidatively damaged DNA, MutM (Fpg), from an extreme thermophile, Thermus thermophilus HB8. EMBO J., 19, 3857–3869.

