## Supplementary files for "Catalytic and Structural Insights into Neil3-Dependent Unhooking of Endogenous Abasic DNA Crosslink"

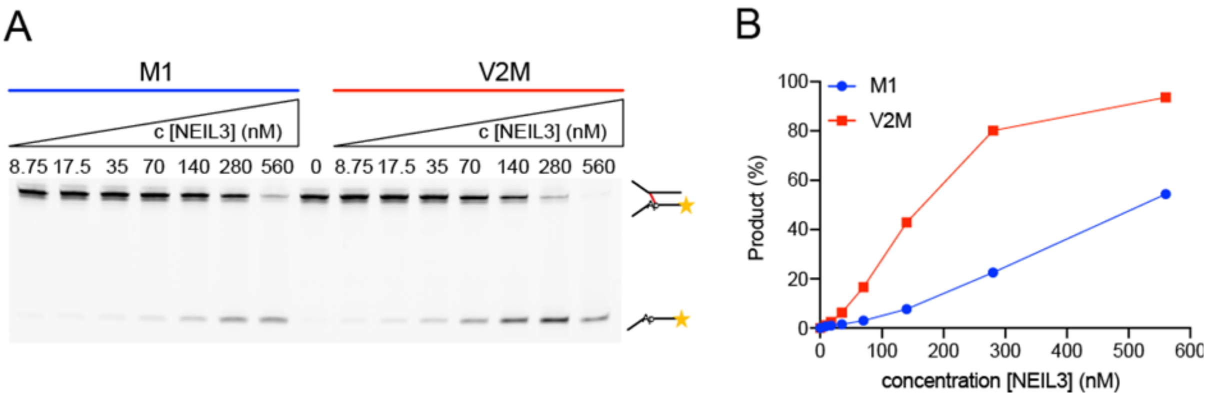


***Supplementary Figure 1:*** **Differential processing of Ap-ICL by NEI domain variants.** *Titration-dependent cleavage of an Ap-ICL containing fork substrate by M1 and V2M NEI (A). Gel analyses were quantified, and cleavage efficiencies are presented as a bar chart (B).*

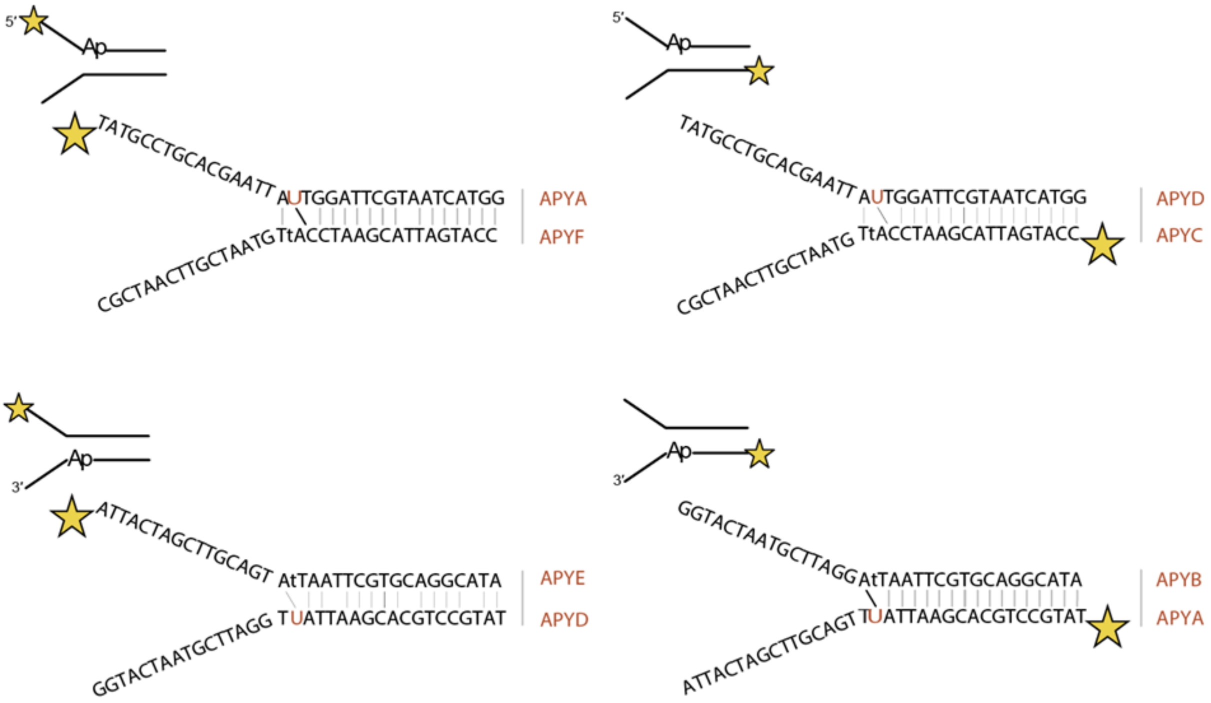


| | |
|---|---|
| APYA | [ATTO488]TATGCCTGCACGAATTAUTGGATTCGTAATCATGG |
| APYB | ATTACTAGCTTGCAGTAtTAATTCGTGCAGGCATA |
| APYC | [ATTO488]CCATGATTACGAATCCAtTGTAATCGTTCAATCGC |
| APYD | TATGCCTGCACGAATTAUTGGATTCGTAATCATGG |
| APYE | [ATTO488]ATTACTAGCTTGCAGTAtTAATTCGTGCAGGCATA |
| APYF | CCATGATTACGAATCCAtTGTAATCGTTCAATCGC |

***Supplementary Figure 2:*** **Design of Ap-ICL and Ap-site containing DNA fork substrates.** *The scheme illustrates DNA substrates in which an Ap site is generated from uracil and subsequently forms an Ap-ICL at different positions within the substrate. The star denotes the ATTO488 fluorescent label. The table below the scheme lists the sequences of the DNA oligonucleotides used.*
